# Dogs’ behaviour is more similar to that of children than to that of cats in a prosocial problem situation

**DOI:** 10.1101/2025.04.27.650646

**Authors:** Melitta Csepregi, Anna Ágnes Moravcsik, Ádám Miklósi, Márta Gácsi

## Abstract

Investigating prosociality within a comparative framework is crucial for understanding the evolutionary origins and emergence of prosocial behaviours, often considered uniquely human. We compared 16-24-month-old children and two distantly related domesticated species, untrained companion dogs and cats—both living in the human niche among relatively similar conditions but characterised by different ecological, evolutionary, and developmental backgrounds. We assessed subjects’ spontaneous behaviour in a natural helping situation, where a familiar human caregiver (parent/owner) searched for an object hidden by the experimenter in the subject’s presence without requesting help. We measured orientation and object-related behaviours, distinguishing those explainable by stimulus enhancement (approaching/manipulating) from the ones most probably explainable by prosocial behaviours (showing/fetching). To control for motivation, we added a trial where subjects’ favourite treat/toy was hidden.

All three species showed similar levels of attention towards the caregiver/object. In line with our hypotheses, children and dogs displayed similar levels of object-related behaviour in the test trials, including those explainable also by stimulus enhancement and those likely indicative of prosociality. In contrast, cats only displayed showing (gaze alternation), and even this occurred with a significantly lower probability. Importantly, in the motivational trial, no species differences were observed in any variable, indicating that cats could also be involved in the problem situation if it was their own interest. Our findings suggest that domestication and close social contact with humans do not, by themselves, induce a tendency towards spontaneous, human-like prosociality. The similar results in dogs and children can be explained by dogs’ social/cooperative nature inherited from their ancestor and the specific selection effects that occurred during their domestication. These factors may have contributed to dogs’ exceptional cooperativity with humans, even favouring the development of spontaneous interspecific prosocial tendencies.

## 1. Introduction

Prosocial behaviours benefit another individual, sometimes without any apparent benefit to the actor (Jensen et al., 2014). The behaviour should be intentional, not an accidental side effect of a different behaviour, and voluntary (i.e., the actor can act otherwise or not at all; Kopp et al., 2024). Helping, a type of prosocial behaviour, is defined as “*assisting another individual in achieving an action-based goal upon the cognitive appraisal of the specific situation or needs of others*” (Kopp et al., 2024). Humans show prosocial actions from early life. Infants as young as 14–18 months engage in instrumental helping tasks, such as assisting someone in retrieving out-of-reach objects (Warneken and Tomasello, 2009). By 2–4 years, children consistently help even unfamiliar individuals with diverse tasks (Dunfield and Kuhlmeier, 2013).

It is a long-standing debate whether humans are the only species capable of behaving prosocially towards non-kin (Jensen, 2016; Tomasello et al., 2005). Great apes, particularly chimpanzees (*Pan troglodytes*), are prime candidates due to their social nature and phylogenetic closeness to humans. While there is some anecdotal and experimental data for such behaviours, overall, evidence for prosociality in primates is inconsistent (Jensen, 2016; Kopp et al., 2024). Recently, prosocial behaviours were investigated in various non-human animal species across other taxonomic groups, with varied and often ambiguous results (Charron et al., 2024; Marshall-Pescini et al., 2016).

Considering their function in the animals’ natural environment, prosocial behaviours occur mostly among conspecifics. However, prosociality was also observed in some interspecific contexts. For example, chimpanzees (Hepach et al., 2020; Warneken and Tomasello, 2006) and hand-raised wolves (Range et al., 2019) were reported to show some forms of prosocial behaviour towards a human partner in specific contexts. However, these contexts often involve excessive pre-training and rewards for the actor, rendering it impossible to evaluate spontaneous prosociality.

Based on their social nature, domestication history, and developmental environment, dogs (*Canis familiaris*) are ideal candidates to show spontaneous prosocial behaviours, especially in an interspecific context (Miklósi et al., 2004). We aimed to explore prosocial behaviour from an evolutionary perspective by comparing three distantly related species—companion dogs, cats, and children—all living in the human niche under relatively similar conditions.

During early domestication, dogs were selected for increased sociality (Driscoll et al., 2009; Trut et al., 2009), which led to high dependency on humans and the development of a human-analogous attachment behaviour between dogs and their owners (Gácsi et al., 2013; Kovács et al., 2018; Topál et al., 2005). Recently, certain dogs have been selectively bred to cooperate with humans, such as in herding or hunting. These breeds are now also suited for new roles as assistance or service dogs (Reeve et al., 2021). Experiments conducted with dogs across various—sometimes ecologically irrelevant—contexts, failed to find consistent evidence of prosociality in dog-dog (e.g., Dale et al., 2019b, 2019a) and dog-human interactions (e.g., Bräuer et al., 2013; Kaminski et al., 2011; Quervel-Chaumette et al., 2016; Range et al., 2019).

Although cats (*Felis catus*) and dogs fulfil similar roles in our lives as companion animals, occupying the same niche, there are considerable differences in their ecological and domestication history and in certain keeping conditions. Domestic cats’ ancestors, a non-social species, formed a loose, mutually beneficial relationship with humans by hunting rodents around human settlements (Driscoll et al., 2009). Thus, cats were not specifically domesticated for integration into the human social environment. Cats reportedly share specific socio-cognitive skills with dogs yet fall short in others when compared to them (Pongrácz and Lugosi, 2024). Cats might exhibit empathic behaviours towards humans, although they seem less likely to show behaviours that can be interpreted as empathetic concern when compared to dogs (Hiestand, 2023). There is a lack of available data on prosocial behaviour in companion cats.

We investigated whether children, dogs, and cats show spontaneous prosocial behaviours in a problem situation where a familiar human partner (owner/parent) searches for a hidden object. We designed a relatively natural test situation requiring no prior training and added a motivational trial when it was the subjects’ own interest to participate in the interaction.

Our general hypothesis was that living in close social contact with humans and undergoing domestication, in general, does not provide an adequate background for a tendency for spontaneous, human-like prosociality in cats. Instead, an inherent social nature and/or specific selective pressures—such as those seen in dogs—are necessary. Therefore, we assumed that, like children but unlike cats, dogs would spontaneously indicate the location of a hidden, irrelevant object to their familiar caregiver without receiving a direct reward. We expected similar attention levels across species due to the inclusion of highly social cats. Confusion, indicating engagement but difficulty inferring the situation, was predicted to be less frequent in cats, with no difference between dogs and children. Object-related behaviours were selected to distinguish stimulus enhancement from prosociality, leading us to expect fewer such behaviours in cats but similar levels in dogs and children.

## 2. Methods

### 2.1. Ethical Note

Before the experiment, parents and owners (hereinafter referred to as caregivers) were informed about the detailed circumstances and goals of the study and signed an informed consent form. Participation was voluntary, and the data obtained were used solely for scientific purposes. Caregivers could stop participating in the experiment at any time. Data collection and all experimental protocols were approved for children by the United Ethical Review Committee for Research in Psychology (EPKEB; Permission 2023-12) and for dogs and cats by the University Institutional Animal Care and Use Committee (Ref. No. ELTE-AWC-009/2023). All methods were carried out under relevant guidelines and regulations, including the Declaration of Helsinki. Personally identifiable data were treated confidentially and stored separately from the rest of the research data per applicable data protection laws.

### 2.2. Subjects

Subjects were selected based on an application questionnaire completed by their caregivers. We included: 1) dogs that were not highly trained (i.e., participated in intermediate obedience training at most), had no specific training for fetching, and spent at least half their time indoors for a valid comparison with the other groups; 2) cats that spent at least half their time indoors and were not fearful of strangers; 3) children who could not yet speak in complete sentences, ensuring that they could not ask about what their parent was searching for, making them comparable to the other groups.

Companion dogs (N = 40), companion cats (N = 27) and human children (N = 20) participated in the test with their caregivers. For detailed information regarding subjects’ age and sex, see Table 1. Dogs belonged to 27 different breeds, in addition to 8 mongrels. Cats did not belong to specific breeds, except 1 Bengal, 1 British shorthair, 1 Maine coon, 1 Scottish straight, and 1 Siamese. We balanced the sex and age of the subjects across the groups as much as possible.

**Table 1.**
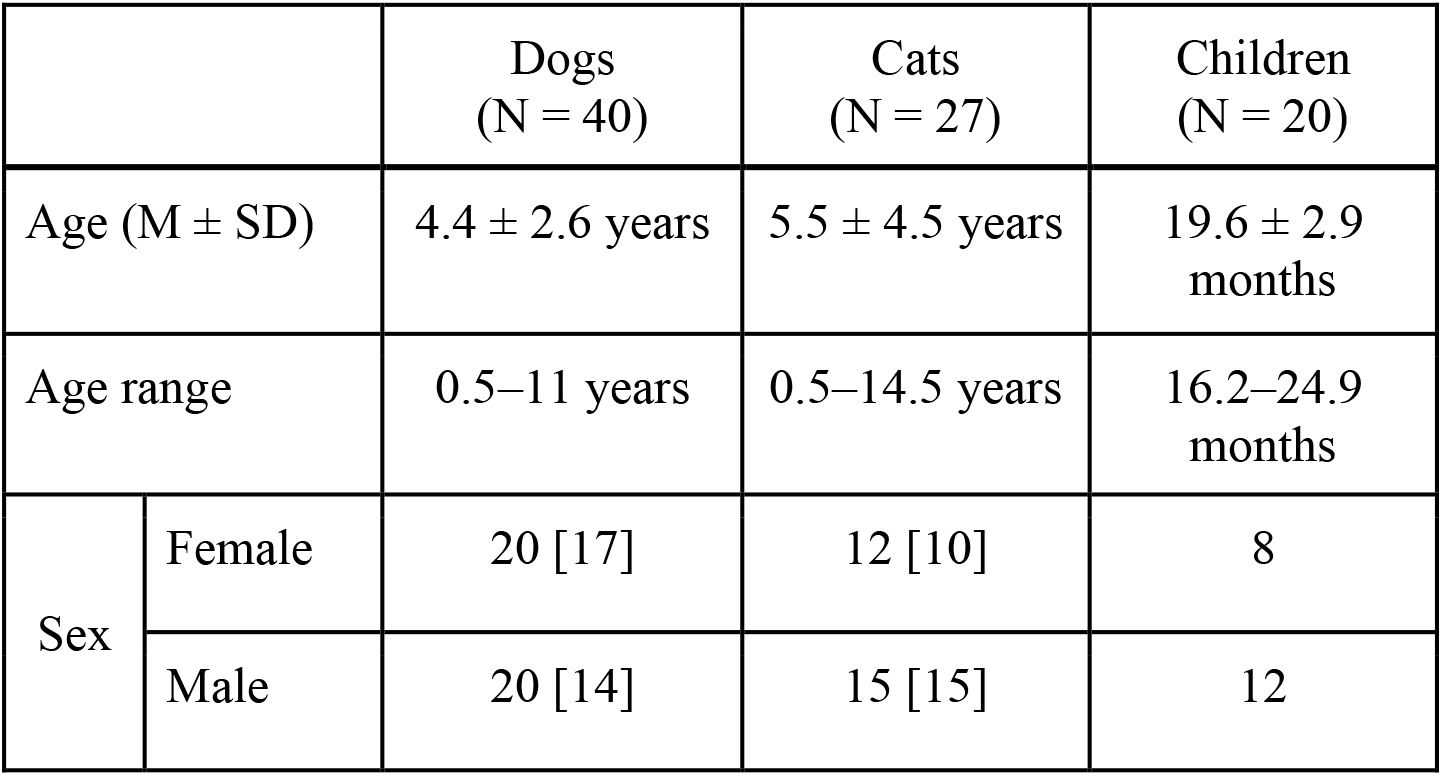
Detailed information regarding subjects’ age and sex, with the number of spayed/neutered individuals in brackets.

We excluded two dogs, five cats, and one child from the analyses, although they all passed the habituation procedure. The reason behind their exclusion was either that the caregiver did not correctly follow the protocol (1 dog and 1 child) or that the subject excluded themselves from the test situation after a while, i.e., spent at least half of a trial hiding or standing/sitting in the door while orienting at the door (1 dog and 5 cats). For more detailed information on the subjects included in the analyses, see Table S1 of the Supplementary material.

Based on the questionnaire data provided by the owners for the subjects included in the analyses, there was no difference between dogs and cats in the following aspects: 1) the amount of time the owner spent actively engaging with the animal, 2) the amount of time the animal spent inside the house, 3) training levels (nothing/basic vs. tricks/tasks), and 4) their affinity to spontaneously fetch objects.

### 2.3. Questionnaires

Caregivers completed an online questionnaire to apply for the experiment. The questionnaire consisted of basic demographic questions about the caregivers and the subjects. In the case of dogs and cats, we assessed the animals’ keeping conditions, training background (with special attention to fetching-related experiences), and openness towards strangers. In the case of the children, we assessed the child’s language skills and attitudes towards strangers. For the questionnaire, see Section S2 of the Supplementary material.

### 2.4. Procedure

The test was always conducted at a predetermined time of the day at the caregiver’s home, which was agreed upon in advance with the respective caregiver (C). We chose a time when the subject was relatively active but not used to engage in any other specific routine activity (e.g., feeding or walking time). Since cats generally appear more challenging to motivate during behaviour tests when compared to dogs (e.g., Kraus et al., 2014; Salamon et al., 2023), we requested cat owners to refrain from feeding their cats for approximately three hours before the test. The experimenter was always the same 25-year-old female.

Before the experiment started, the experimenter (E) explained the procedure in detail to C. During the habituation phase, E attempted to interact with the subject to help them become accustomed to E’s presence. This habituation phase lasted 10 minutes, but it could be concluded earlier if the subject showed no signs of fear towards E. To be considered successfully habituated to E’s presence, dogs and cats had to fulfil at least two of the following three criteria: play with E, accept E’s treats, allow E to pet them for at least 15 seconds. Similarly, children were considered successfully habituated if they fulfilled at least two criteria: accept an object from E or voluntarily give an object to E, voluntarily establish physical contact with E, approach and smile at E.

During the habituation phase, E carefully showed the subject any new objects (e.g., hand-held camera, test object), allowing them to investigate the items until they lost interest. This procedure aimed to ensure that subjects would not display object-related behaviours driven by their own interest. The habituation and the experiment were conducted in a closed room, usually a bedroom or living room, relatively free from distractions (e.g., toys, food).

#### 2.4.1. Warm-up trial

The purpose of this trial was to familiarise C and the subject with the test situation. Initially, C and E handled the object (a brand new, clean dishwashing sponge with a scrubby underside, size 8×4.5×2.5 cm) for 15 seconds. Afterwards, E asked C to turn away while she put the object away in a visible place. C was asked to look for the object for a few seconds (visibly looking around and searching while saying, “Oh no, I can’t find it! What should I do now?” repeatedly). Once C found the object, he/she reacted enthusiastically and excitedly.

#### 2.4.2. Testing phase

##### Test trial 1

C and E enthusiastically interacted with the object (which was no longer of interest or relevance to the subject). If the subject attempted to take the object away, both C and E ignored the subject and prevented access to the object. After approximately 30 seconds, E instructed C to turn away and face a corner of the room. E captured the subject’s attention (e.g., by calling their name) and waited for eye contact. If the subject was frustrated or distracted by something, E waited until they returned to a neutral state and tried to establish eye contact again. Once eye contact was established, E moved the object to an unreachable location, such as on top of a shelf, and concealed it with a small towel. Following this, E waited 5 seconds or until the subject lost interest in the hidden object and moved away.

Afterwards, E instructed C to turn back, and C began searching for the object while expressing dismay with repeated utterances (the same as in the warm-up trial). C did not talk directly to the subject during this phase. However, C had to look at the subject occasionally while searching to give the subject a chance to establish eye contact. Meanwhile, E stood passively in the corner of the room, measuring the elapsed time using a handwatch.

We observed whether the subject spontaneously indicated the location of the hidden object to C (i.e., approached or pointed at the hiding spot, alternated their gaze between C and the object, manipulated the object if possible). If they did so, C could obtain the object. Alternatively, if the subject did not do so within 30 seconds, E told C about the whereabouts of the hidden object. When C acquired the object (either with the help of the subject or E), he/she responded enthusiastically upon receiving it.

##### Test trial 2

Regardless of how C found the object the previous time, the procedure of Test trial 1 was repeated. However, in this case, E hid the same object in a new location where the subject could see it but could not reach it (e.g., on the edge of a shelf without covering it with the towel).

##### Test trial 3

The same procedure was repeated regardless of how C found the object the previous time. However, in this case, E hid the same object in a new location where the subject could see and reach it (e.g., on the floor), thus gradually making the solution easier.

##### Motivational trial

The procedure of Test trial 2 was repeated, but in this trial, E hid an object that was important or interesting to the subject (e.g., a favourite toy or food) instead of the sponge. This trial evaluated the subject’s motivation and ability to exhibit the measured behaviours. This trial was always conducted last to prevent subjects from associating the problem situation with (self-)rewards, thus allowing us to assess their spontaneous prosociality in the test trials.

### 2.5. Behavioural Variables

Test sessions were videotaped, and the recordings were coded using BORIS version 8.20.4 (Friard and Gamba, 2016).

We assessed the subjects’ spontaneous behaviour across three main aspects (similarly to Csepregi and Gácsi, 2023): 1) attention directed towards the relevant aspects of the environment (owner/parent, hidden object), 2) confusion-related behaviours suggesting interest in the partner’s emotional state but difficulty comprehending the task, and 3) object-related behaviours that could be interpreted as prosocial behaviour (Miklósi et al., 2000; Pongrácz and Lugosi, 2024; Virányi et al., 2006; Zhang et al., 2021). The logic we used to assess subjects’ behaviour in the test is illustrated in Figure 1.

**Figure 1.**
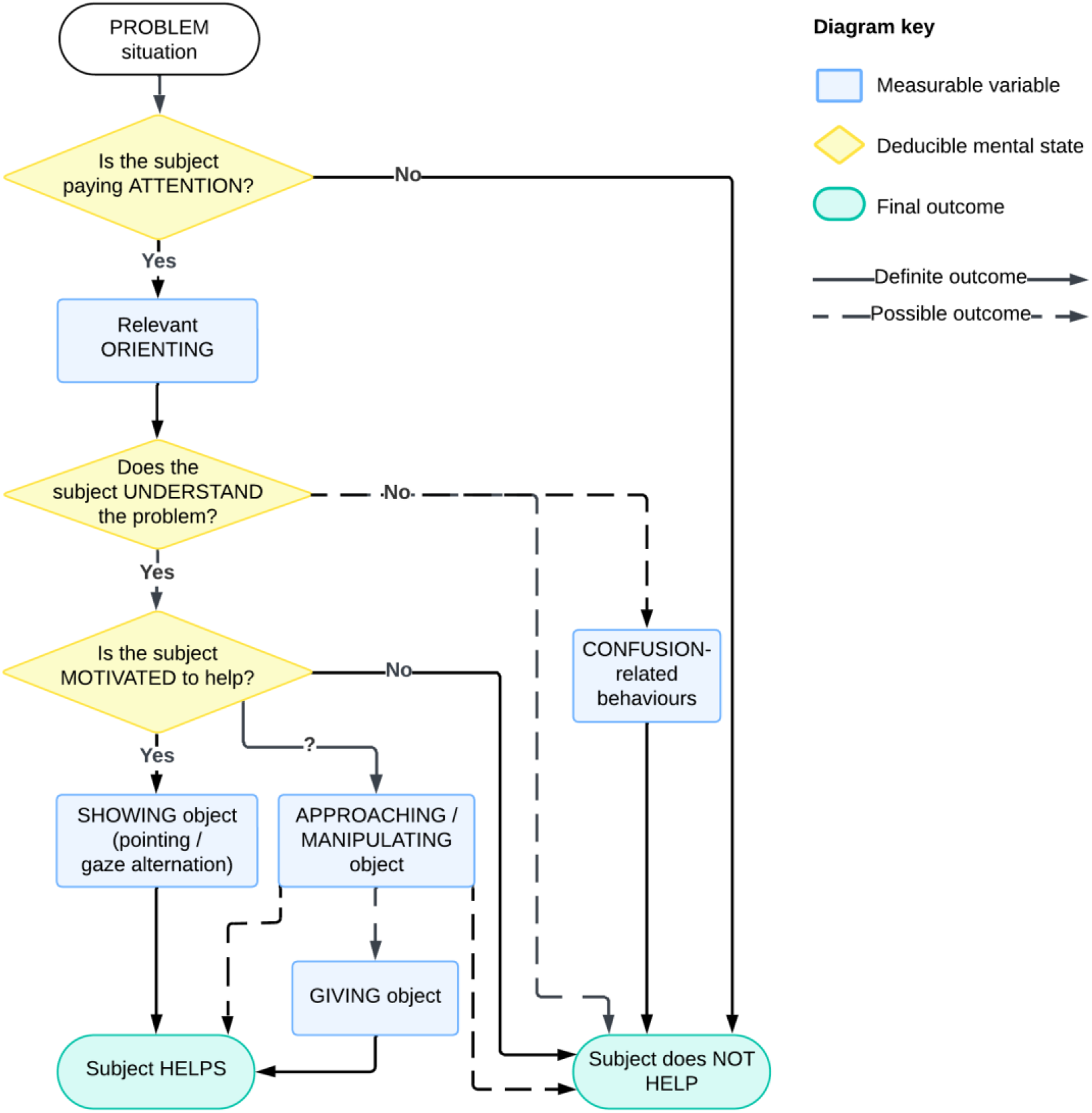
The decision model we propose to interpret paradigms investigating prosocial behaviour (the actual behaviours must be matched to the given task). The model allows us to deduce the underlying mental process behind each step. While the term “understanding” is admittedly difficult to define and often interchanged by other synonyms (e.g., infer, comprehend, appraise, assess), we used this word for simplicity, as others also did in previous studies on non-human animals (e.g., Kaminski et al., 2011; Marshall-Pescini et al., 2016; Quervel-Chaumette et al., 2016).

For a list of the measured behavioural variables corresponding to each of these components and their definitions, refer to Table 2. In each trial, the given behaviours were coded during the caregiver’s search for the object (i.e., from the moment the caregiver turned around until 30 seconds passed or until the subject indicated the object’s correct location). Examples of the coded behavioural variables are illustrated in Figure 2.

**Table 2.** The coded behavioural variables and their detailed definitions. The underlined components and variables were modelled in the statistical analysis. The test trials included trials 1–3.

| **Variable name** | **Definition** | **Measure** | **Trials when coded** |
| --- | --- | --- | --- |
| Relevant orientation at object or caregiver | The amount of time the subject spends orienting towards the hidden object or the caregiver | duration (s) | Test trials, Motivational trial |
| Vocalising while close to caregiver | Any kind of vocalisation (e.g., barking, meowing, babbling), except purring, while the subject is in the caregiver's proximity (in reaching distance) and does not show gaze alternations. Coded only before the very first occurrence of showing/giving (prosocial) behaviour | yes/no | Test trials |
| Approaching | Approaching the object or the surface on which the object is located within the subject's body length in the case of dogs, and cats and within reaching distance in the case of children, while orienting at the object | yes/no | Test trials, Motivational trial |
| Showing | Gazing at the caregiver is directly followed by a gaze at the object and back within 2 seconds or vice versa (gaze alternation); or, in the case of children, pointing with the hand in the direction of the hidden object | yes/no |  |
| Manipulating | Mouthing, pawing, touching or holding the object | yes/no | Test trial 3 |
| Giving | Giving the object to the caregiver | yes/no | Test trial 3 |

**Figure 2.**
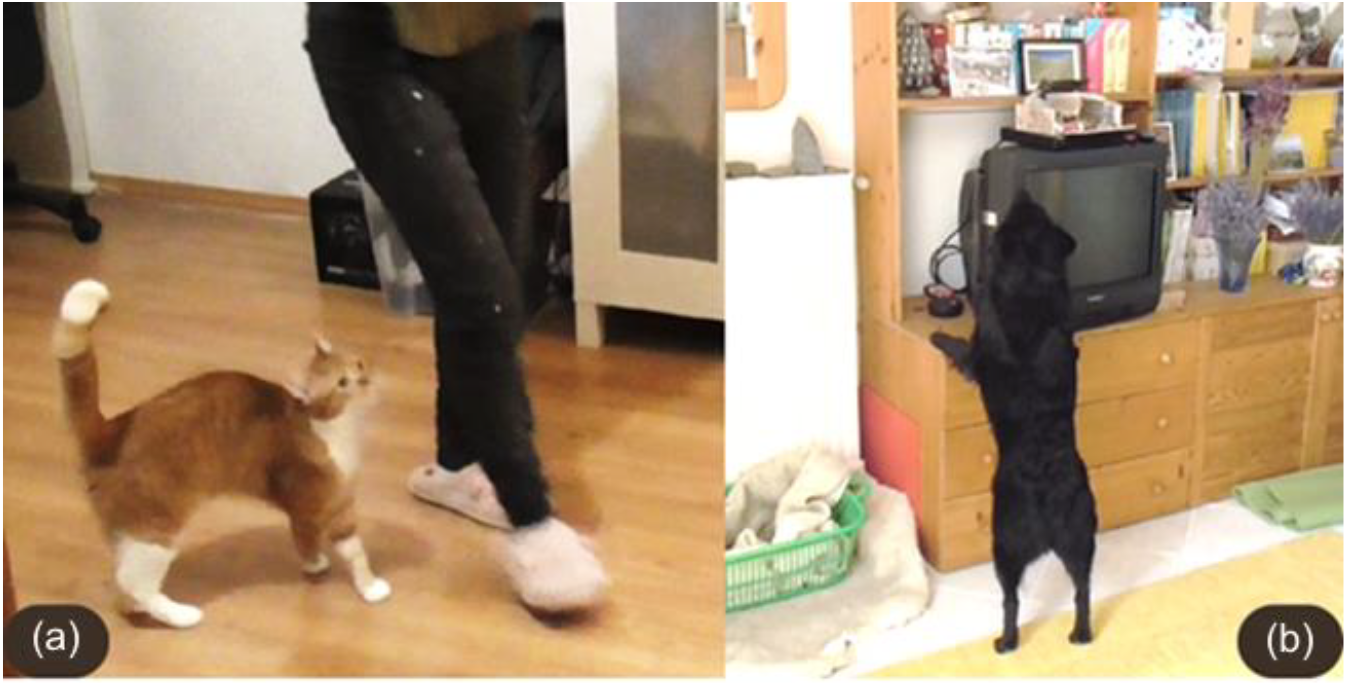
Examples of the following behavioural variables: a) orienting at the caregiver, and b) approaching the object.

### 2.6. Statistical Analysis

The inter-rater reliability was assessed on a subsample where all four coders independently coded 20% of the subjects. Cohen’s kappa values revealed moderate to strong reliability (M ± SD = 0.708 ± 0.160 for all coded variables). Due to variations in trial duration, time percentages were used to measure relevant orientation.

Statistical analyses were performed in software R (v4.4.1; R Core Team, 2023). The Shapiro-Wilk test and normal probability plots (‘qqplotr’ package) showed that none of the continuous behavioural variables were normally distributed. We used generalised linear mixed models (GLMM) for analysis (‘lme4’ and ‘glmmTMB’ packages). Count models were checked for zero inflation (‘performance’ package).

#### Test trials

To model relevant orientation, we used a GLMM, which included species and trial number as fixed factors, and subject ID as a random effect. The Gamma family function was used.

A negative binomial GLMM was used to model vocalising while close to the caregiver. Species and trial number were included as fixed factors, and subject ID was included as a random effect.

To allow for clear interpretations, we treated all object-related variables separately. Separate binomial GLMMs were used for approaching and showing behaviour, with species and trial number as fixed factors and subject ID as a random effect. Cats did not show approaching in the test trials; thus, they were excluded from the model. In the case of children, we used the Kruskal-Wallis test to assess whether age affected the displayed type of showing behaviour (gaze alternation/pointing).

For all models, post hoc pairwise comparisons between factors were conducted using the Tukey test with Benjamini–Hochberg adjustment (‘emmeans’ package).

Section S3 of the Supplementary Material presents the GLMM results for fixed effects and the post hoc pairwise comparisons to enhance readability.

For object manipulation, which was only possible in Test trial 3, cats had to be excluded from the analysis because this behaviour did not occur in their case. Thus, Pearson’s chi-squared test was used to assess differences between dogs and children. Fisher’s exact test was used to compare the likelihood of giving the object to the caregiver after manipulation (for dogs and children).

#### Motivational trial

The Kruskal–Wallis test was used to assess differences between species in relevant orientation. Vocalising while close to the caregiver could not be analysed statistically, as neither dogs nor cats showed such behaviour in the motivational trial. For approaching and showing, Pearson’s chi-squared test was used. The Benjamini-Hochberg correction was applied to control for false positives due to multiple comparisons.

In this trial, the object (the subject’s favourite food/toy) was hidden in a visible but unreachable location, similar to the protocol of Test trial 2. Thus, object manipulation could not occur.

#### Behaviour patterns

Cluster analysis was performed to identify patterns in attention and object-related behaviours across the test trials. For the attention variable, we calculated the mean duration of relevant orientation across all three trials for each subject. We created a derived score for each subject to analyse object-related behaviours based on their exhibited behaviours. The scoring system was as follows: a score of 2 was assigned if the subject showed the object (via gaze alternation or pointing) or gave it to the caregiver in at least one of the three trials; a score of 1 was assigned for approaching or manipulating the object (without showing or giving it) in at least one of the three trials; and a score of 0 was given if neither behaviour occurred.

After scaling the variables, we determined the optimal number of clusters using the elbow method, which suggested *k* = 4 clusters based on the within-cluster sum of squares. Clusters were formed using the k-means method, with the algorithm run 100 times to ensure stability. The Kruskal-Wallis test was then used to compare the clusters based on attention, followed by Dunn’s tests for pairwise comparisons with p-values adjusted using the Benjamini–Hochberg method. Fisher’s exact test was used to compare the clusters based on object-related behaviour scores, followed by pairwise test comparisons with p-values adjusted using the Benjamini– Hochberg method (‘pairwiseNominalIndependence’ function). Detailed results of the pairwise comparisons are provided in Section S4 of the Supplementary Material to enhance readability.

## 3. Results

### Test trials

In the case of relevant orientation, no significant difference was found between species or test trials (Fig. 3, Supplementary Table S2).

**Figure 3.**
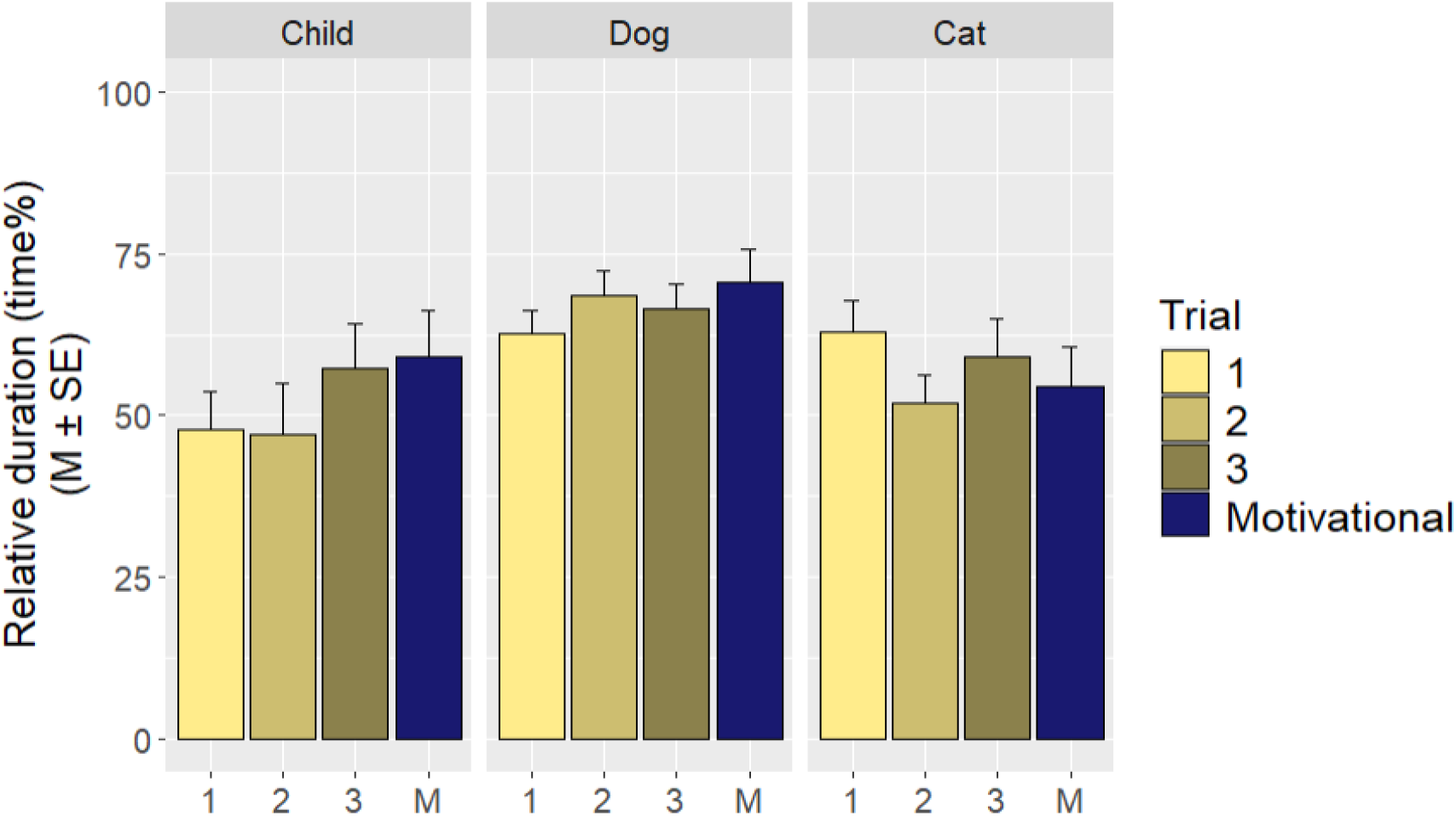
The relative duration of relevant orientation (i.e., directed at the caregiver or the hidden object) across species and trials.

There was a significant difference between species in vocalising while close to the caregiver: children were more likely to show such behaviour than dogs or cats. No difference was found between test trials (Fig. 4, Supplementary Table S3-4).

**Figure 4.**
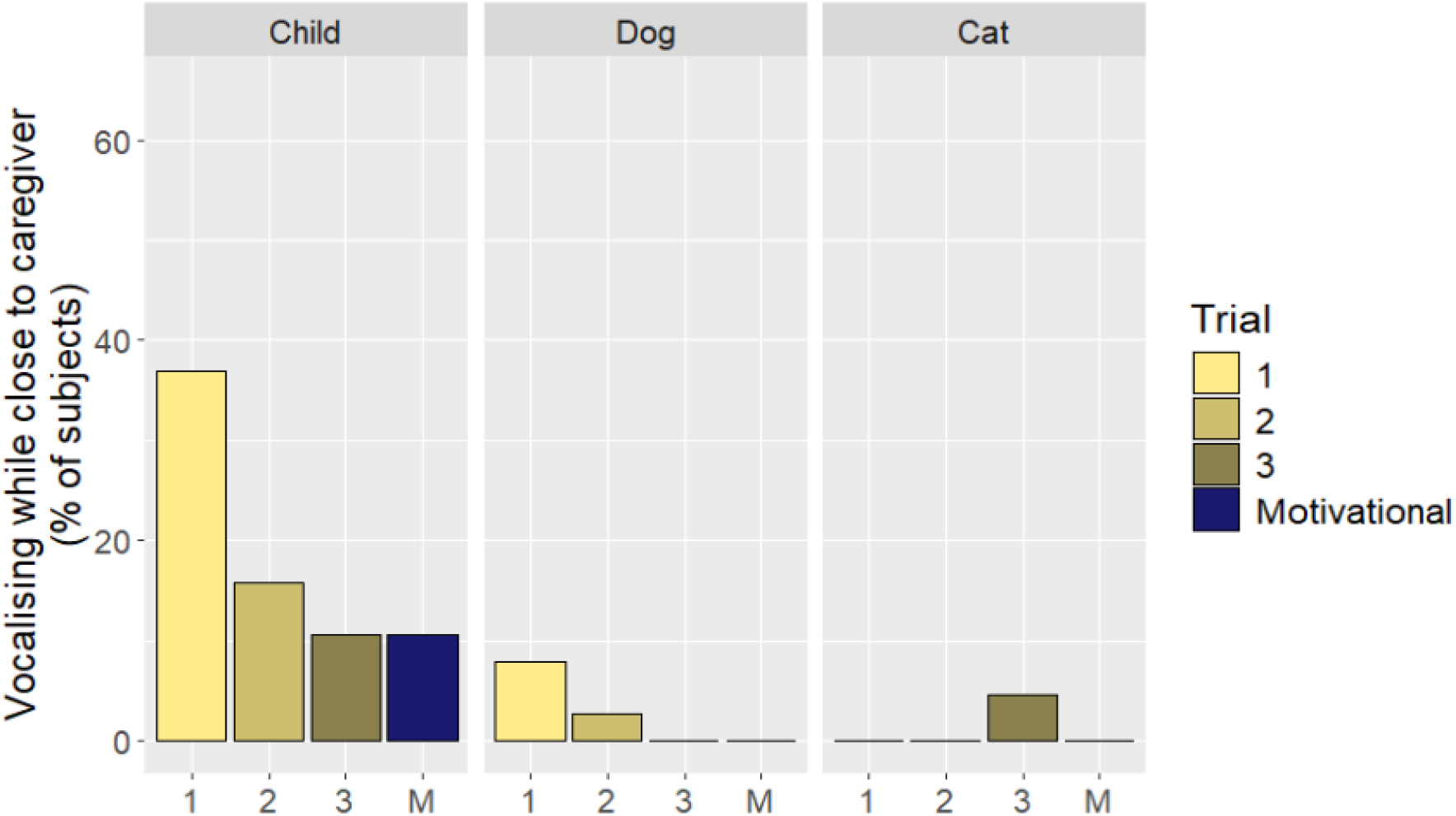
The occurrence of vocalising while close to the caregiver across species and trials (coded only before the very first occurrence of prosocial behaviour).

No cats displayed approaching behaviour; thus, they were not included in the analysis. No significant difference was observed between dogs and children. Approaching was significantly more likely to occur in Test trials 2 and 3 than in Test trial 1. Similarly, it was more likely to occur in Test trial 3 than in Test trial 1 (Fig. 5a, Supplementary Tables S5-6).

**Figure 5.**
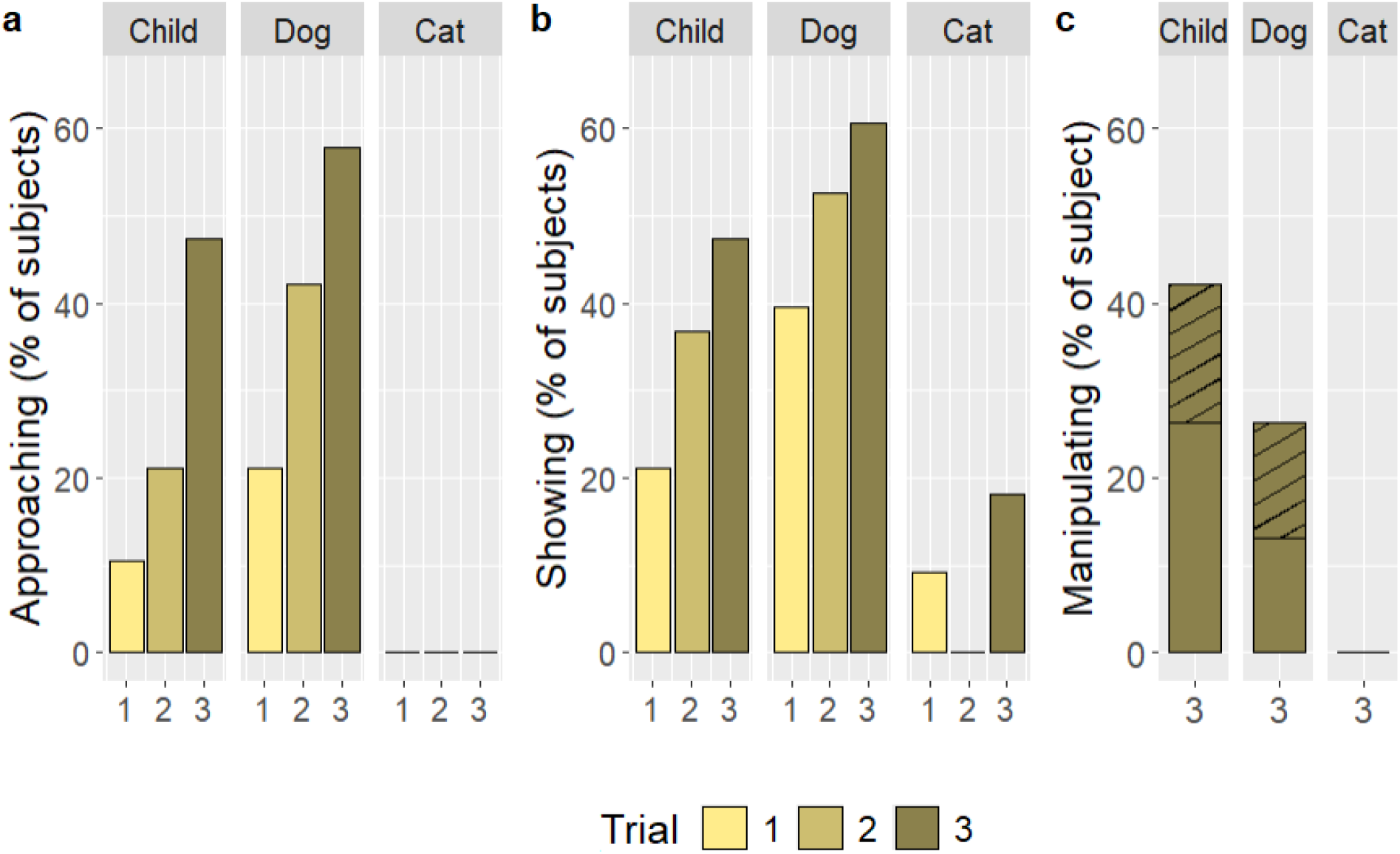
The occurrence of a) approaching, b) showing, and c) manipulating the hidden object across species in the test trials. Object manipulation was possible in Trial 3 only; the stripes indicate subjects that gave the object to the caregiver.

Significant differences were found between species in the showing behaviour: dogs and children were more likely to show the hidden object than cats. Showing was more likely to occur in Test trial 3 than in Test trial 1 (Fig. 5b, Supplementary Tables S7-9). In the case of children, the different means of showing were distributed in the test trials as follows: 15.8% used pointing only, 15.8% used gaze alternation only, 31.6% used both behaviours, and 36.8% used none. Age did not affect children’s displayed type of showing behaviour (H_3_ = 4.425, *P* = 0.219).

Cats were excluded from the statistical test of object manipulation as none of them touched the sponge. There was no significant difference between dogs and children (χ2_1_ = 1.462, *P* = 0.227). Also, no difference was found between dogs and children in whether they gave the object to the caregiver (*P* = 0.637) (Fig. 5c).

For an overview of object-related behaviours of the three species across the test trials, see Figure 6.

**Figure 6.**
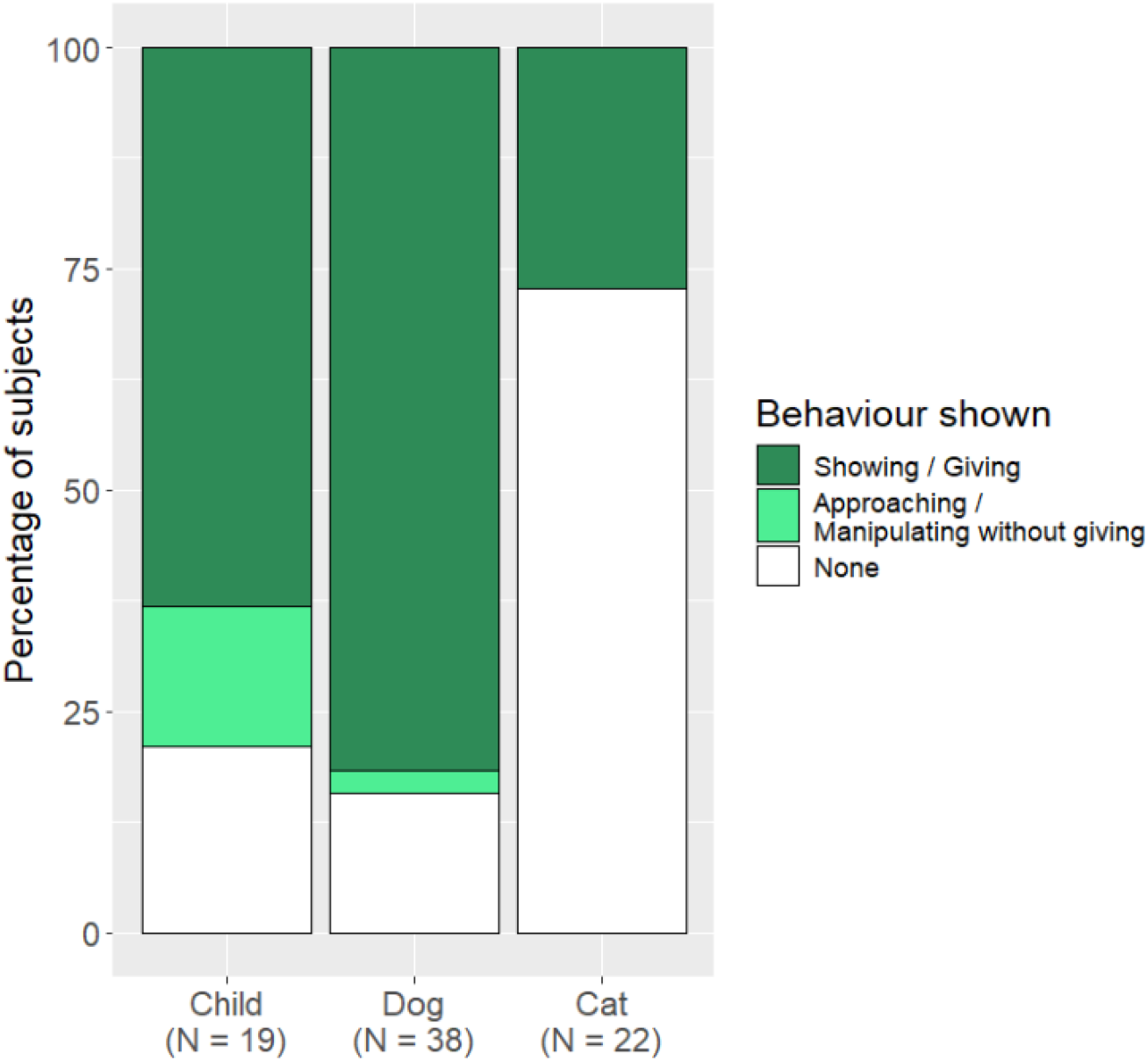
Object-related behaviours across species in the test trials. The dark green represents subjects displaying prosocial behaviours (showing, giving the object to C). The light green indicates subjects exhibiting behaviours that could be attributed to either stimulus enhancement or prosociality (approaching the object, manipulating it without giving it to C). Each subject is represented only once: if a subject exhibited behaviours from both categories across the test trials, they were included exclusively in the prosocial category.

### Motivational trial

In the motivational trial, when the subjects’ favourite food/toy was hidden (based on the protocol of Test trial 2), there was no difference between species in terms of relevant orientation (H_2_ = 5.609, *P* = 0.061) (Fig. 3).

There was no significant difference between species regarding approaching behaviour (χ2_2_ = 6.760, *P* = 0.035; non-significant after Benjamini-Hochberg correction) (Fig. 7a).

**Figure 7.**
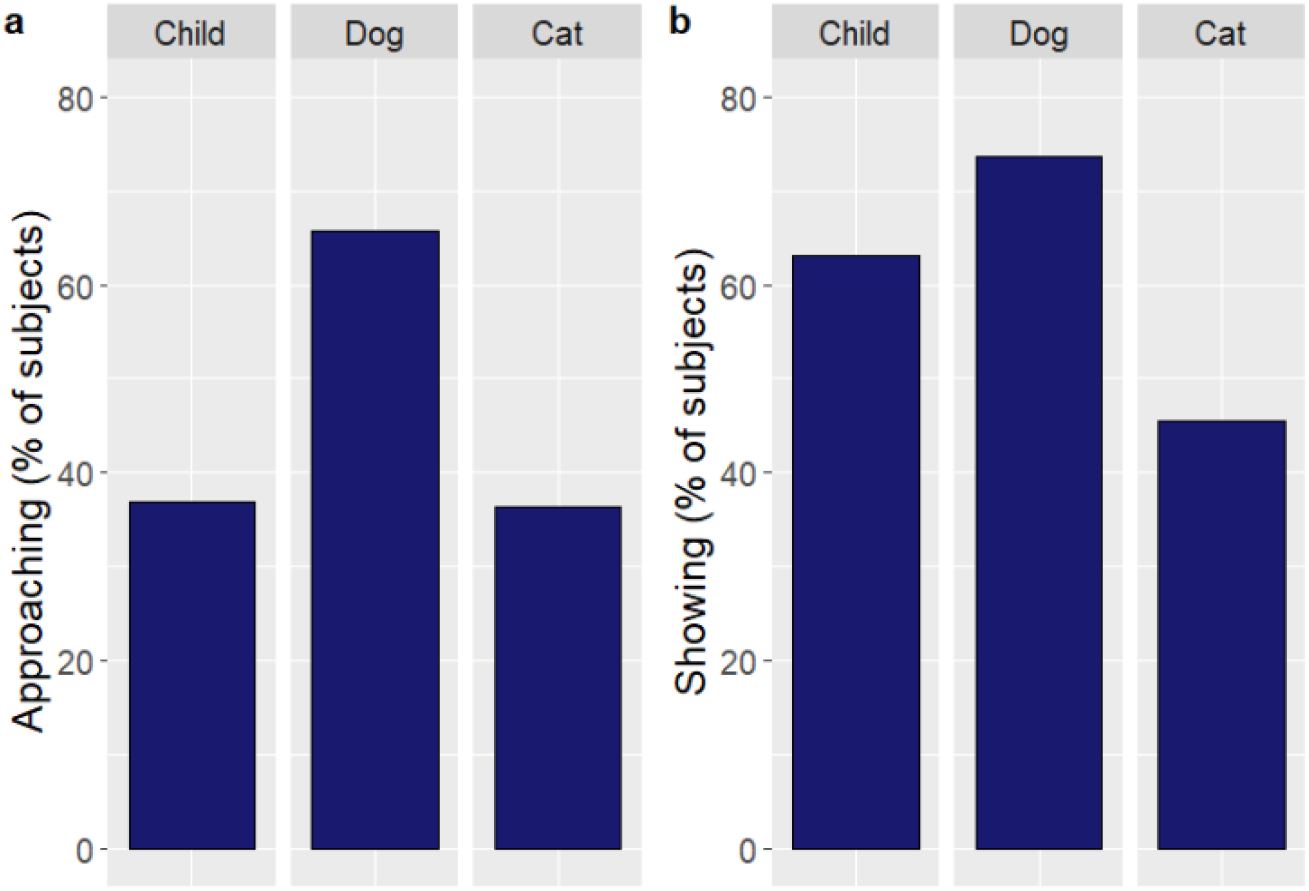
The occurrence of object a) approaching and b) showing behaviours across species in the motivational trial. In this trial, the object (subject’s favourite food/toy) was hidden in a visible but unreachable location, similar to the protocol of Test trial 2.

No significant difference was found between species for showing behaviour (χ2_2_ = 4.779, *P* = 0.092) (Fig. 7b).

### Behaviour patterns

Figure 8 shows the cluster analysis results. Table 3 provides the mean values for attention, the medians of object-related behaviours in each of the four clusters, and the distribution of each species across clusters.

**Table 3.** Descriptive data of the four clusters.

| **Cluster** | **Mean attention**<br>**(time %)** | **Object-related**<br>**behaviour score**<br>**(0-2) (median)** | **Distribution**<br>**of species**<br>**between clusters** |
| --- | --- | --- | --- |
| 1 | 19.94 | 0 | Child: 21.1%<br>Dog: 2.6%<br>Cat: 18.2% |
| 2 | 52.84 | 2 | Child: 52.6%<br>Dog: 36.8%<br>Cat: 18.2% |
| 3 | 65.17 | 0 | Child: 5.2%<br>Dog: 15.8%<br>Cat: 54.5% |
| 4 | 80.14 | 2 | Child: 21.1%<br>Dog: 44.7%<br>Cat: 9.1% |

**Figure 8.**
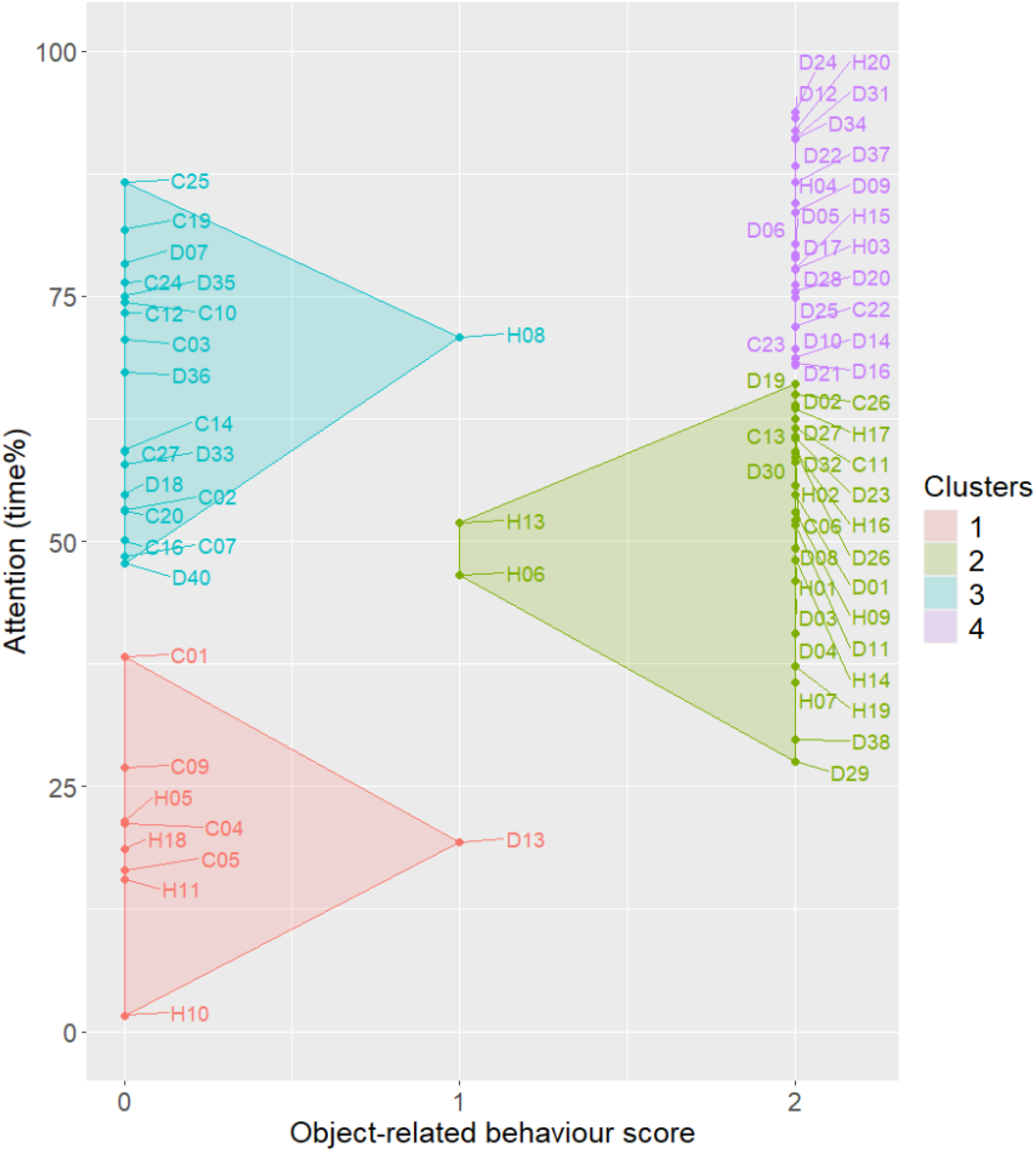
The four clusters represent the patterns in subjects’ attention and object-related behaviours across the test trials. (H: human, C: cat, D: dog)

For attention, the mean time percentage spent orienting at the caregiver/object differed significantly among the four clusters (H_3_ = 57.799, *P* < 0.001). Post hoc tests revealed significant differences between all clusters (Supplementary Table S10). Object-related behaviours also varied significantly among the clusters (*P* < 0.001), with significant differences observed between all clusters except for 1 and 3, and 2 and 4, respectively (Supplementary Table S11).

## 4. Discussion

Our study aimed to explore prosocial behaviour from an evolutionary perspective by comparing the behaviour of children, untrained companion dogs, and companion cats—three species sharing similar environments but differing in ecological and evolutionary backgrounds. Cats were significantly less likely to exhibit spontaneous object-related behaviours than dogs and children when it did not directly benefit them. The similar object-related behaviours of children and dogs suggest that, in certain contexts, dogs tend to spontaneously help their human caregivers to the same degree as 16-24-month-old children, even without a direct reward.

In line with our hypothesis, cats spent no less time orienting towards the object/caregiver than dogs and children, indicating that they were interested in the situation regardless of their later involvement in problem-solving. Though (Fugazza et al., 2023) found that dog puppies paid more attention to the human demonstration of a simple action than kittens in a laboratory test, our result aligns with the findings of (Bogese et al., 2024), who reported similar gaze patterns in cats and dogs during everyday interactions with the owner. We explain our result primarily by the preselection of our cat subjects for increased social openness towards humans so that they would be more motivated to participate in the test (Salamon et al., 2023).

Confusion-related behaviours were more likely to occur in children than in dogs or cats. This difference likely stems from the nature of our measured behavioural variable (vocalising while close to caregiver), as children were generally more vocal than the other two species. Additionally, few dogs were attentive without displaying object-related behaviours, which may explain the few confusion-related behaviours. This aligns with our decision model (Fig. 1), where we suggested that confusion-related behaviour should only occur if a subject is attentive and motivated to help (or at least sensitive to the caregiver’s emotional state) but does not understand the situation (which would manifest in object-related behaviours).

We considered different forms of object-related actions to distinguish between prosocial motivation and alternative explanations. Approaching and manipulating the hidden object may have resulted from either stimulus enhancement, where subjects perceive the object as important based on caregivers’ contextual cues, or an intrinsic motivation to help, as these behaviours could serve as a form of showing, thus qualifying as prosocial behaviour. Cats’ lack of approaching behaviour likely reflects a lower tendency to adjust their behaviour to the owner’s emotional state or to synchronise their movement with their owners’ (Duranton et al., 2017) rather than a consequence of being too fearful of getting involved in the situation (since our subjects were preselected and habituated). In the motivational trial, where engagement aligned with their own interests (i.e., approaching behaviour could be interpreted as requesting rather than helping), cats approached the hidden object at similar rates to dogs and children. This suggests they assessed the situation but engaged only when it benefitted themselves, consistent with their relative independence from humans (Pongrácz and Lugosi, 2024) and lack of selection for cooperative behaviours during domestication (Driscoll et al., 2009). Additionally, cats’ ancestors were solitary animals—although nowadays, domestic cats are considered “socially flexible”, dynamically adapting to different forms of social life, including living in groups with conspecifics and/or humans (Crowley et al., 2020). However, due to the fixed order of the trials, it cannot be excluded that cats inferred the situation much slower than dogs and children, only during the final motivational trial.

Object manipulation occurred at similar rates in children and dogs but was never observed in cats. While the habituation phase was carried out to exclude/reduce spontaneous object interest, caregiver actions during the test may have reignited it in the case of dogs and children. Thus, manipulating the object could have resulted from stimulus enhancement. However, giving the object to the caregiver must reflect a prosocial intent, which also occurred with a similar likelihood in dogs and children but never in cats. Notably, dogs exhibited this behaviour despite lacking prior training in object-fetching tasks.

We considered showing behaviour a key indicator of prosociality. Spontaneous showing behaviour was significantly more frequent in dogs and children than in cats, but importantly, it occurred also in the case of cats. Notably, during the test trials, subjects received no direct reward for signalling the object’s location, ruling out the expectation of individual benefit. This species-specific difference emerged only in test trials, where helping the caregiver provided no direct advantage, but disappeared in the motivational trial. Notably, the overall proportion of children displaying object-related behaviours (Fig. 6) was similar to previous studies on “altruistic helping” in children of similar age (Warneken et al., 2007; Warneken and Tomasello, 2006).

As the test trials progressed, the likelihood of approaching and showing the hidden object increased. This suggests that subjects engaged more with the situation over time as its complexity gradually decreased, and they likely gained a better assessment of the problem. Those individuals who were attentive but neither helped nor showed confusion may have understood the problem but simply chose not to assist (see Fig. 1).

Although we carefully balanced our sample on certain relevant factors (such as training experience) as much as possible, some developmental factors might have contributed to the behavioural differences between cats and dogs. It cannot be ruled out that dogs engage in more diverse types of social interactions with their owners; for example, they are disciplined more often than cats. However, cats in our study may also have had an advantage over dogs, as only carefully preselected, non-fearful cats were included, some of which were trained for tasks and tricks—traits not necessarily representative of companion cats in general. Conversely, we excluded highly trained dogs, representing only the “lower end” of training experience. The difficulties in balancing all aspects are mostly due to the inherent differences between the two species, which do not allow for testing subjects with completely identical individual development (e.g., most cats cannot be taken for walks or tested in a lab (Uccheddu et al., 2022).

We argue that the occurrence of prosocial behaviour requires understanding the social partner’s problem, which may not have been consistently achieved in prior studies (e.g., Kaminski et al., 2011; Quervel-Chaumette et al., 2016). Despite partly similar experimental contexts (indicating the location of a hidden object to the owner), we successfully demonstrated prosocial behaviour in dogs, contrasting with (Kaminski et al., 2011). In our study, the owner and experimenter jointly manipulated the object with evident interest before every trial, likely capturing the dog’s attention. Additionally, we hid only one object at a time, aiding the dogs’ assessment of their owner’s search target. By contrast, Kaminski et al. had owners perform functional tasks (e.g., cutting or stapling paper), possibly failing to highlight the object’s relevance. We argue that understanding the problem and its emotional aspects is a prerequisite for prosocial behaviour (Fig. 1). Our results reinforce that experimental contexts must be cognitively and socially accessible when assessing prosociality.

A critical distinction in our study is that subjects that helped did not merely engage in a shared activity with the caregiver, as seen in studies where dogs interpreted the searching as a social game (Topál et al., 2006). If dogs’ participation had been driven solely by behavioural synchronisation (Kubinyi et al., 2003), they would not have indicated the location of the object or handed it over to the owner.

Notably, it is uncertain whether dogs recognised the owner’s lack of knowledge regarding the hiding place. Nevertheless, their ability to infer the owner’s searching behaviour (and negative emotional state) during the search phase was sufficient to elicit helping responses. Thus, while we think having presented evidence for prosocial behaviour in dogs and children, this experiment does not provide a clear insight into the cognitive processes controlling this behaviour.

Some previous studies have failed to identify either interspecific (Quervel-Chaumette et al., 2016) or intraspecific (Dale et al., 2019b) prosociality in dogs. Moreover, many relied on excessive pre-training, food rewards, and/or food-sharing contexts (which introduces competition) (e.g., Melis et al., 2006; Satoh et al., 2021; Schwab et al., 2012). An additional key advantage of our method was the absence of direct rewards during test trials, allowing us to observe truly spontaneous prosocial behaviours.

The cluster analysis results (Fig. 8) support our decision model (Fig. 1). Cluster 1 comprised individuals with minimal attention and object-related behaviours, showing no prosociality. Cluster 2, mainly consisting of children, displayed moderate attention and consistent object-related behaviours, with prevalent prosocial responses. This suggests that our attention variable (orientation) may not have been entirely suitable for measuring attention in children, which may be more precisely assessed via eye-tracking (de Jong et al., 2016) or more focused video recordings (Choudhury and Gorman, 2000). Cluster 3, predominantly cats, included attentive subjects who either did not comprehend the situation or were not motivated to help. Cluster 4, the most homogeneous one, contained individuals displaying high attention and object-related behaviours, typically dogs. These subjects completed all steps of the proposed decision model (Fig. 1).

Our results show that dogs, similar to 16-24-month-old children, are likely to engage in spontaneous interspecific prosocial behaviours with their caregivers in an instrumental helping task. However, in cats, domestication, living in the human niche, and forming close social bonds with humans did not commonly induce human-like prosociality. Based on our decision model, this could be explained either by the cats’ less understanding of the problem situation (owner in need) or by being less motivated to help the owner.

The proper assessment of the test situation requires various skills. Although cats were reported to show comparable skills to that of dogs in some interspecific communicative tasks (Miklósi et al., 2005; Pongrácz et al., 2019), in a recent direct comparative study, dogs outperformed cats both in their testability and in relying on human distal pointing gestures (Salamon et al., 2023).

Furthermore, Miklósi et al. (2005) found that dogs looked at the human and back to the hidden food when unable to reach the reward, while cats persevered longer, trying to solve the problem individually, rarely looking at the human. Similarly, only dog puppies but not kittens showed a spontaneous tendency to match their behaviour actions to human demonstrations in the absence of food rewards (Fugazza et al., 2023).

Dogs’ strong motivation to help is presumably related to the unique dog-owner relationship (Topál and Gácsi, 2012), supported by a recent comparative study according to which even a social domesticated species, the miniature pig (*Sus scrofa domesticus*), does not show dog-like attachment behaviour toward their owners (Gábor et al., 2024).

Our findings highlight the role of species-specific evolutionary factors in shaping social behaviours. The dog’s unique status as a cooperative companion animal likely stems from multiple contributing factors: its social inheritance from a pack-hunting ancestor, a unique domestication process inducing dependence on owners, and a tendency for social/cooperative interactions with humans. These factors, alongside some unavoidable differences in keeping conditions (which are not independent of these evolutionary factors), likely explain why dogs exhibit social skills more akin to children’s than another domesticated companion animal species, the cat.

## Acknowledgements

We thank all parents and owners for participating in the study and Sarolt Kinga Gintner and Nóra Sziklai for their valuable help in recruiting subjects and coding the video recordings. This research was funded by the HUN-REN–ELTE Comparative Ethology Research Group (F01/031), and the Ministry of Innovation and Technology of Hungary from the National Research, Development, and Innovation Office—NKFIH (K132372).

