## Supplementary material for "Dogs’ behaviour is more similar to that of children than to that of cats in a prosocial problem situation"

**Section S1**

*Table S1.* Demographic data on the subjects included in the statistical analyses.

| **ID** | **Species** | **Age at testing (months)** | **Sex** | **Breed** |
| --- | --- | --- | --- | --- |
| D01 | Dog | 12.0 | Neutered male | mongrel |
| D02 | Dog | 89.2 | Neutered male | mongrel |
| D03 | Dog | 82.8 | Neutered male | Labrador retriever |
| D04 | Dog | 8.6 | Intact female | Braque d'Auvergne |
| D05 | Dog | 15.6 | Neutered male | miniature poodle |
| D06 | Dog | 72.0 | Neutered female | mudi |
| D07 | Dog | 97.6 | Neutered female | Alaskan malamute |
| D08 | Dog | 60.3 | Neutered female | standard schnauzer |
| D09 | Dog | 33.7 | Neutered female | Old English sheepdog |
| D10 | Dog | 71.6 | Neutered female | golden retriever |
| D11 | Dog | 54.9 | Intact male | pumi |
| D12 | Dog | 60.0 | Neutered male | mongrel |
| D13 | Dog | 132.0 | Neutered male | mongrel |
| D14 | Dog | 38.7 | Neutered male | dachshund (short haired) |
| D16 | Dog | 7.7 | Intact female | Hungarian vizsla |
| D17 | Dog | 24.6 | Neutered female | dachshund (wire haired) |
| D18 | Dog | 90.8 | Neutered female | Havanese |
| D19 | Dog | 12.7 | Neutered male | mongrel |
| D20 | Dog | 31.2 | Neutered female | fox terrier (wire haired) |
| D21 | Dog | 31.2 | Neutered female | border collie |
| D22 | Dog | 99.0 | Neutered male | Jack Russell terrier |
| D23 | Dog | 11.4 | Intact male | Jack Russell terrier |
| D24 | Dog | 36.6 | Neutered male | Lagorai shepherd |
| D25 | Dog | 37.8 | Neutered male | Havanese |
| D26 | Dog | 97.9 | Neutered female | American pitbull terrier |
| D27 | Dog | 47.5 | Intact male | Scottish terrier |
| D28 | Dog | 61.0 | Intact female | English cocker spaniel |
| D29 | Dog | 59.6 | Neutered female | English cocker spaniel |
| D30 | Dog | 40.2 | Neutered male | mongrel |
| D31 | Dog | 46.0 | Neutered female | mongrel |
| D32 | Dog | 66.6 | Intact male | Labrador retriever |
| D33 | Dog | 44.7 | Neutered male | golden retriever |
| D34 | Dog | 90.6 | Neutered male | komondor |
| D35 | Dog | 60.0 | Neutered female | mongrel |
| D36 | Dog | 70.7 | Neutered female | fox terrier (short haired) |
| D37 | Dog | 30.4 | Intact male | whippet |
| D38 | Dog | 6.0 | Intact male | whippet |
| D40 | Dog | 25.2 | Neutered female | shiba inu |
| C01 | Cat | 8.0 | Intact female | housecat |
| C02 | Cat | 166.6 | Neutered female | housecat |
| C03 | Cat | 102.7 | Neutered male | housecat |
| C04 | Cat | 109.9 | Neutered male | housecat |
| C05 | Cat | 110.6 | Neutered male | Siamese |
| C06 | Cat | 10.8 | Neutered female | housecat |
| C07 | Cat | 132.0 | Neutered female | housecat |
| C09 | Cat | 57.9 | Neutered male | housecat |
| C10 | Cat | 117.3 | Neutered female | housecat |
| C11 | Cat | 53.9 | Neutered male | housecat |
| C12 | Cat | 173.8 | Neutered female | housecat |
| C13 | Cat | 19.4 | Neutered female | Scottish straight |
| C14 | Cat | 36.0 | Neutered female | housecat |
| C16 | Cat | 6.5 | Intact female | housecat |
| C19 | Cat | 154.1 | Neutered male | housecat |
| C20 | Cat | 12.,0 | Neutered male | housecat |
| C22 | Cat | 11.0 | Neutered male | housecat |
| C23 | Cat | 31.3 | Neutered male | housecat |
| C24 | Cat | 24.0 | Neutered male | Maine coon |
| C25 | Cat | 21.0 | Neutered female | housecat |
| C26 | Cat | 38.2 | Neutered male | housecat |
| C27 | Cat | 13.1 | Neutered male | housecat |
| H01 | Child | 16.3 | Female | - |
| H02 | Child | 16.5 | Female | - |
| H03 | Child | 19.4 | Male | - |
| H04 | Child | 16.8 | Male | - |
| H05 | Child | 19.7 | Male | - |
| H06 | Child | 23.6 | Female | - |
| H07 | Child | 19.8 | Male | - |
| H08 | Child | 19.2 | Female | - |
| H09 | Child | 19.1 | Male | - |
| H10 | Child | 18.8 | Female | - |
| H11 | Child | 18.1 | Male | - |
| H13 | Child | 17.2 | Male | - |
| H14 | Child | 18.4 | Female | - |
| H15 | Child | 16.2 | Male | - |
| H16 | Child | 16.6 | Male | - |
| H17 | Child | 19.1 | Male | - |
| H18 | Child | 23.0 | Male | - |
| H19 | Child | 24.9 | Female | - |
| H20 | Child | 24.2 | Female | - |

**Section S2**

The following questionnaires were used to assess subjects’ demographic data and background. Bulleted points indicate available categories provided for each question.

*Questionnaire for dog owners*

1. Owner’s name
2. Email address, used only for the communication necessary for organising the test and is not passed on to third parties.
3. The dog’s name
4. The dog’s sex
   - Male
   - Female
5. Is the dog spayed/neutered?
   - Yes
   - No
6. The dog’s date of birth in YYYY.MM.DD format. If you do not know your dog’s exact birth date, please, enter the following numbers: ʺ1212.12.12.ʺ and go to the next question where you can enter the estimated age of the dog.
7. Your dog’s estimated date of birth. If you do not know exactly when your dog was born, please estimate its age.
8. The dog’s breed
9. How open is your dog towards strangers? Score on a Likert scale from 1 (“My dog is shy and distrustful towards strangers”) to 5 (“My dog is very open towards anyone”).
10. Keeping conditions
    - My dog spends at least half of its time indoors.
    - My dog spends approx. equal time indoors and outdoors.
    - My dog spends at least half of its time outdoors.
11. On average, how much time do you actively spend with your dog daily?
    - Less than 1 hour
    - 1-2 hours
    - 3-4 hours
    - More than 4 hours
12. Training experience
    - I only taught my dog basic necessities
    - My dog participated in basic obedience training
    - My dog participated in something else (other than basic obedience training), but does not have a certificate
    - My dog has a certificate or is actively competing in sports/training
    - My dog is an assistance dog (therapy dog, guide dog, etc.)
13. Does your dog like to carry/fetch objects (regardless of training)? Score on a Likert scale from 1 (“Not at all”) to 5 (“Very much”).
14. How experienced is your dog in fetching tasks?
    - My dog is not experienced in fetching tasks at all
    - If I throw its toy, my dog usually brings it back
    - My dog can fetch simple objects (not only toys/sticks/dummies) when it is given a command
    - My dog can fetch any object, keep it in its mouth and put it down when it is given a command (might have difficulties with metal objects)
    - My dog can put any object in/on top of other objects and can be commanded to bring an object from one person to another

*Questionnaire for cat owners*

1. Owner’s name
2. Email address, used only for the communication necessary for organising the test and is not passed on to third parties.
3. The cat’s name
4. The cat’s sex
   - Male
   - Female
5. Is the cat spayed/neutered?
   - Yes
   - No
6. The cat’s date of birth in YYYY.MM.DD format. If you do not know your cat’s exact birth date, please, enter the following numbers: ʺ1212.12.12.ʺ and go to the next question where you can enter the estimated age of the cat.
7. Your cat’s estimated date of birth. If you do not know exactly when your cat was born, please estimate its age.
8. The cat’s breed
9. How open is your cat towards strangers? Score on a Likert scale from 1 (“My cat is shy and distrustful towards strangers”) to 5 (“My cat is very open towards anyone”).
10. Keeping conditions
    - My cat spends at least half of its time indoors.
    - My cat spends approx. equal time indoors and outdoors.
    - My cat spends at least half of its time outdoors.
11. On average, how much time do you actively spend with your cat daily?
    - Less than 1 hour
    - 1-2 hours
    - 3-4 hours
    - More than 4 hours
12. Training experience
    - My cat is not trained at all
    - My cat is trained only for some basic necessities (e.g., no biting, no jumping on the kitchen counter)
    - My cat can perform some tricks and tasks as well
13. Does your cat like to carry/fetch objects (regardless of training)? Score on a Likert scale from 1 (“Not at all”) to 5 (“Very much”).
14. Have you trained your cat for fetching tasks?
    - Yes
    - No

*Questionnaire for parents*

1. Parent’s name
2. Email address, used only for the communication necessary for organising the test and is not passed on to third parties.
3. The child’s first name
4. The child’s sex
   - Male
   - Female
5. The child’s date of birth
6. How open is your child towards strangers? Score on a Likert scale from 1 (“My child is shy and distrustful of strangers”) to 5 (“My child is very open towards anyone”).
7. What is your child’s level of speech development?
   - Only babbles
   - Uses one or two words
   - In about half of the cases uses gestures and sounds, in the other half expresses himself/herself with words
   - Usually says what he/she wants and often asks questions with words
   - Uses phrases and shorter sentences
   - Other…
8. How interested is your child in new, unfamiliar objects? Score on a Likert scale from 1 (“Not interested”) to 5 (“Very interested”).
9. On average, how much time do you actively spend with your child daily? (Excluding feeding and hygiene activities.)
   - Less than 1 hour
   - 1-2 hours
   - 3-4 hours
   - More than 4 hours

**Section S3**

Detailed description of the GLMMs reported in the Results for each behavioural variable, along with the results of the models for fixed effects, and post hoc pairwise comparisons (with Benjamini-Hochberg adjustment). In the GLMMs, cats are treated as a baseline for species, and Trial 1 for trials.

*Table S2.* Relevant orientation in Trials 1-3.

|  | Estimate | Std. Error | t value | Pr(>\|z\|) |
| --- | --- | --- | --- | --- |
| (Intercept) | 0.018 | 0.002 | 9.331 | < 0.001 |
| Species_dog | -0.002 | 0.002 | -1.148 | 0.251 |
| Species_child | 0.002 | 0.003 | 1.038 | 0.299 |
| Trial_2 | 0.000 | 0.002 | 0.055 | 0.956 |
| Trial_3 | -0.001 | 0.002 | -0.463 | 0.644 |

*Table S3.* Vocalising while close to caretaker in Trials 1-3.

|  | Estimate | Std. Error | z value | Pr(>\|z\|) |
| --- | --- | --- | --- | --- |
| (Intercept) | -4.111 | 1.154 | -3.563 | < 0.001 |
| Species_dog | 0.786 | 1.171 | 0.672 | 0.502 |
| Species_child | 2.625 | 1.111 | 2.362 | 0.018 |
| Trial_2 | -0.917 | 0.592 | -1.549 | 0.121 |
| Trial_3 | -1.204 | 0.658 | -1.829 | 0.067 |

*Table S4.* The pairwise comparisons of vocalising while close to caretaker across species.

| Contrast | Odds ratio | Std. Error | z ratio | p value |
| --- | --- | --- | --- | --- |
| cat / dog | 0.455 | 0.533 | -0.672 | 0.502 |
| cat / child | 0.073 | 0.081 | -2.362 | 0.027 |
| dog / child | 0.159 | 0.110 | -2.647 | 0.024 |

*Table S5.* Approaching object in Trials 1-3 (for dogs and children only, with dogs treated as a baseline for species).

|  | Estimate | Std. Error | z value | Pr(>\|z) |
| --- | --- | --- | --- | --- |
| (Intercept) | -1.777 | 0.493 | -3.598 | < 0.001 |
| Species_child | -0.649 | 0.541 | -1.200 | 0.230 |
| Trial_2 | 1.275 | 0.518 | 2.465 | 0.014 |
| Trial_3 | 2.327 | 0.569 | 4.091 | < 0.001 |

*Table S6.* The pairwise comparisons of approaching behaviour across test trials.

| Contrast | Odds ratio | Std. Error | z ratio | p value |
| --- | --- | --- | --- | --- |
| Trial_1 / Trial_2 | 0.280 | 0.145 | -2.465 | 0.021 |
| Trial_1 / Trial_3 | 0.098 | 0.055 | -4.091 | < 0.001 |
| Trial_2 / Trial_3 | 0.349 | 0.162 | -2.274 | 0.023 |

*Table S7.* Showing object in Trials 1-3.

|  | Estimate | Std. Error | z value | Pr(>\|z\|) |
| --- | --- | --- | --- | --- |
| (Intercept) | -3.339 | 0.651 | -5.132 | < 0.001 |
| Species_dog | 2.820 | 0.625 | 4.512 | < 0.001 |
| Species_child | 1.990 | 0.662 | 3.003 | 0.003 |
| Trial_2 | 0.500 | 0.413 | 1.211 | 0.226 |
| Trial_3 | 1.190 | 0.423 | 2.813 | 0.005 |

*Table S8.* The pairwise comparisons of showing behaviour across species.

| Contrast | Odds ratio | Std. Error | z ratio | p value |
| --- | --- | --- | --- | --- |
| cat / dog | 0.060 | 0.037 | -4.512 | < 0.001 |
| cat / child | 0.137 | 0.091 | -3.003 | 0.004 |
| dog / child | 2.295 | 1.110 | 1.713 | 0.087 |

*Table S9.* The pairwise comparisons of showing behaviour across test trials.

| Contrast | Odds ratio | Std. Error | z ratio | p value |
| --- | --- | --- | --- | --- |
| Trial_1 / Trial_2 | 0.607 | 0.250 | -1.211 | 0.226 |
| Trial_1 / Trial_3 | 0.305 | 0.129 | -2.813 | 0.015 |
| Trial_2 / Trial_3 | 0.502 | 0.201 | -1.724 | 0.127 |

**Section S4**

Detailed description of the post hoc pairwise comparisons of clusters.

*Table S10.* Attention across clusters in Trials 1-3.

| Comparison | z value | p value (adjusted) |
| --- | --- | --- |
| 1 - 2 | -4.146 | < 0.001 |
| 1 - 3 | -6.557 | < 0.001 |
| 2 - 3 | -2.904 | 0.006 |
| 1 - 4 | -2.589 | 0.001 |
| 2 - 4 | 2.306 | 0.002 |
| 3 - 4 | 5.635 | < 0.001 |

*Table S11.* Object-related behaviours across clusters in Trials 1-3.

| Comparison | p value (adjusted) |
| --- | --- |
| 1 - 2 | < 0.001 |
| 1 - 3 | 1.000 |
| 1 - 4 | < 0.001 |
| 2 - 3 | < 0.001 |
| 2 - 4 | 0.594 |
| 3 - 4 | < 0.001 |
